# A Re-examination of the Role of *AUXIN RESPONSE FACTOR 8* in Developmental Abnormalities Caused by the P1/HC-Pro Viral Suppressor of RNA Silencing

**DOI:** 10.1101/056366

**Authors:** Sizolwenkosi Mlotshwa, Gail J. Pruss, John L. Macarthur, Jason W. Reed, Vicki Vance

**Affiliations:** Department of Biological Sciences, University of South Carolina, Columbia, South Carolina, United States of America; Department of Biology, University of North Carolina, Chapel Hill, North Carolina, United States of America

## Abstract

Plant viral suppressors of RNA silencing induce developmental defects similar to those caused by mutations in genes involved in the microRNA (miRNA) pathway. These abnormalities were originally thought to reflect a pleiotropic impact of silencing suppressors on miRNA control of plant development. However, subsequent work with the P1/HC-Pro potyviral suppressor of silencing showed that global impairment of the miRNA pathway was not responsible for the phenotypical anomalies. More recently, developmental defects caused by a *P1/HC-Pro* transgene under control of the 35S promoter were attributed to moderate upregulation of *AUXIN RESPONSE FACTOR 8 (ARF8)*, a target of miR167. The key piece of evidence in that work was that the developmental defects in the 35S-pro:*P1/HC-Pro* transgenic *Arabidopsis* were greatly alleviated in the F1 progeny of a cross with plants carrying the *arf8-6* mutation. *Arf8-6* is a SALK line T-DNA insertion mutant, a class of mutations prone to inducing transcriptional silencing of transgenes expressed from the 35S promoter. Here we report a re-investigation of the role of *ARF8* in P1/HC-Pro-mediated developmental defects. We characterized the progeny of a cross between our 35S-pro:P1/HC-Pro transgenic *Arabidopsis* line and the same *arf8-6* T-DNA insertion mutant used in the earlier study. The T-DNA mutation had little effect in the F1 generation, but almost all *arf8-6/P1/HC-Pro* progeny had lost the P1/HC-Pro phenotype in the F2 generation. However, this loss of phenotype was not correlated with the number of functional copies of the *ARF8* gene. Instead, it reflected transcriptional silencing of the 35S-pro:P1/HC-*Pro* transgene, as evidenced by a pronounced decrease in P1/HC-Pro mRNA2accompanied by the appearance of 35S promoter siRNAs. Furthermore, *arf8-8*, an independent loss-of-function point mutation, had no detectable effects on P1/HC-Pro phenotype in either the F1 or F2 generations. Together these data argue against the reported role of increased *ARF8* expression in mediating developmental defects in *P1/HC-Pro* transgenic plants.

**Author Summary:** RNA silencing is an important antiviral defense in plants that uses small RNA molecules to target the invading RNA. Plant viruses, however, have countered with proteins that suppress RNA silencing, and one of the best-studied plant viral suppressors of silencing is P1/HC-Pro. When the genetic model plant *Arabidopsis thaliana* is bioengineered to express P1/HC-Pro, the resulting plants display distinct developmental abnormalities. These abnormalities are thought to arise because P1/HC-Pro also interferes with the arm of RNA silencing that uses small RNAs called microRNAs (miRNAs) to regulate expression of the plant's own genes. Earlier work, however, showed that interference with all miRNAs in general could not be responsible for these developmental defects. More recently, it was reported that enhanced expression of a single miRNA-controlled gene, *AUXIN RESPONSE FACTOR 8 (ARF8)*, underlies the developmental defects caused by P1/HC-Pro. However, using the same *ARF8* mutation as that report, as well as a second, independent *ARF8* loss-of function mutation, we now show that mis-regulation of *ARF8* is not responsible for those defects. One or a few key miRNA-controlled factors might, in fact, underlie the developmental defects caused by P1/HC-Pro; however, our results show that *ARF8* is not one of the key factors.

## Introduction

Eukaryotes have evolved an elaborate network of RNA silencing pathways mediated by small regulatory RNAs (for reviews, see [1, 2]). These pathways play two major roles in plants. One is to regulate expression of endogenous genes, and the other is to defend against invading nucleic acids, such as viruses, transposons, and-more recently-transgenes. MicroRNAs (miRNAs) are the small RNAs involved in regulation of endogenous genes and play an important role in development, while small interfering RNAs (siRNAs) function in defense against invading nucleic acids. Both of these classes of small RNA are produced by RNase III-like ribonucleases called DICER-LIKE (DCL) in plants; however, miRNAs and siRNAs differ not only in function, but also in biogenesis. In *Arabidopsis*, DCL1 processes miRNAs from endogenous, imperfectly double-stranded transcripts to produce a duplex containing the miRNA and the opposite strand, the miRNA*. In contrast, siRNAs are processed by DCL2, DCL3, or DCL4 from the perfectly double-stranded RNA that triggers the defensive arm of RNA silencing. MiRNA and siRNA duplexes are methylated by HUA ENHANCER 1 (HEN1), and one strand incorporates into an RNA-induced silencing complex to guide the Argonaute (AGO) silencing effector proteins to complementary target sequences. Thus, partially overlapping biosynthetic pathways lead to two classes of small RNAs with distinct functions.

To counteract antiviral silencing, many plant viruses have evolved proteins that suppress RNA silencing. It was noted early on that transgenic plants expressing P1/HC-Pro, the first discovered plant viral suppressor of silencing [3, 4], exhibited pronounced developmental abnormalities [5], in addition to being suppressed for silencing. These developmental defects were first observed in tobacco [5, 6], but were subsequently characterized in greater detail in *Arabidopsis* [7–9]. In Arabidopsis, P1/HC-Pro-mediated developmental defects include reduced size, serrated leaves, late flowering, and abnormalities in floral morphology resulting in reduced fertility. Some of these developmental abnormalities resemble ones caused by mutations in genes involved in plant miRNA biogenesis and function. In addition, *P1/HC-Pro* transgenic lines display clear defects in both the biogenesis and function of miRNAs. MiRNAs are present at higher levels in the *P1/HC-Pro* lines, but the usually labile miRNA* also accumulates [7, 10]. In addition, although miRNA levels are generally increased, so are their mRNA targets, suggesting a reduction in miRNA function [7, 8]. This correlation of developmental defects with defects in the miRNA pathway led to the logical idea that the abnormalities in morphology are caused by interference with miRNA control of developmental pathways [7, 8].

Following up on this idea, a subsequent study investigated the possibility that ectopic over-expression of *DCL1*, which encodes the Dicer that produces miRNAs in plants, might rescue the defects in the miRNA pathway and thereby alleviate the developmental abnormalities in *P1/HC-Pro* transgenic *Arabidopsis*. Surprisingly, although over-expression of *DCL1* did largely alleviate the developmental defects, it did not correct the P1/HC-Pro-associated defects in the miRNA pathway: levels of miRNAs and their targets were unchanged from those seen in the parental *P1/HC-Pro* line [9]. These data indicate that P1/HC-Pro-mediated defects in development do not result from general impairments in miRNA biogenesis or function. However, because only a few miRNAs and their targets were examined, the study could not rule out the possibility that the developmental defects were due to interference with one or a small set of miRNA-controlled genes. Consistent with that idea, it was later reported that the developmental defects in *P1/HC-Pro* transgenic plants are due to upregulation of *AUXIN RESPONSE FACTOR 8 (ARF8)*, a target of miR167 that plays a key role in auxin signaling and developmental patterning [11].

We were interested in further investigating the role of *ARF8* in P1/HC-Pro-mediated developmental defects because the conclusion that upregulation of *ARF8* is responsible for the defects is in conflict with the earlier work on the effect of overexpression of *DCL1* on P1/HC-Pro phenotype [9]. Importantly, one of the miRNA and target pairs looked at in that work was miR167 and its target, *ARF8*. Increased accumulation of *ARF8* mRNA was unaffected by over-expression of *DCL1* in *P1/HC-Pro* plants, even though the P1/HC-Pro developmental phenotype was largely eliminated. Moreover, the developmental phenotypes of *P1/HC-Pro*-expressing plants differ from those caused by mutation of the *miR167* target site in *ARF8* or the redundantly acting *ARF6*. *P1/HC-Pro*-expressing plants have narrow serrated leaves, narrow sepals and petals, and short stamen filaments [7–9]. In contrast, *mARF8* or *mARF6* plants with mutated *miR167* target sites do not have serrated leaves or narrow flower organs, but do have elongated stamen filaments and swollen (indehiscent) anthers, and are also female sterile due to arrested ovule integument growth [12] Based on those results, upregulation of *ARF8* would not appear to be correlated with or responsible for P1/HC-Pro-mediated developmental defects. To resolve these apparent discrepancies, therefore, we further characterized the effects of *arf8* mutations on the P1/HC-Pro phenotype using the same mutation as in the published work [11] as well as a second, independent mutation.

## Results

### The P1/HC-Pro developmental phenotype is largely retained in the F1 generation of a cross with the *arf8-6* T-DNA insertion mutant

Developmental defects in a *P1/HC-Pro* transgenic *Arabidopsis* line were reported to be greatly alleviated in the F1 progeny of a cross with plants carrying the *arf8-6* mutation [11]. To try to reproduce that result, we used the same *arf8-6* SALK line as in that study and crossed it to our established *P1/HC-Pro* transgenic *Arabidopsis* line [9, 13, 14]. The *P1/HC-Pro* line is hemizygous for the Turnip Mosaic Virus (TuMV) coding region, expresses high levels of the *P1/HC-Pro* mRNA, and has a severe developmental phenotype ([9] and Fig 1). In the F1 generation, about half (124/238) of the progeny of the cross had the P1/HC-Pro phenotype, while about half (114/238) did not. Genotyping 69 of the non-phenotypic plants revealed that none had the *P1/HC-Pro* transgene, and all but three had the *arf8-6* mutation, indicating that absence of P1/HC-Pro phenotype in the F1 progeny population did not reflect any effect of the *arf8-6* mutation but simply reflected absence of the *P1/HC-Pro* transgene. Among 71 F1 plants confirmed to be both hemizygous for *P1/HC-Pro* and heterozygous for *arf8-6*, all had the P1/HC-Pro phenotype. Thus, in our system, hemizygous *arf8-6* does not cause wide-spread, strong reduction in P1/HC-Pro phenotype in the F1 generation. A minority of our P1/HC-Pro phenotypic F1 progeny, however, showed a subtle reduction in developmental abnormalities (Fig 1, F1), suggesting that effects of the *arf8-6* mutation might be better detected in the F2 generation of the cross.

**Fig 1.**
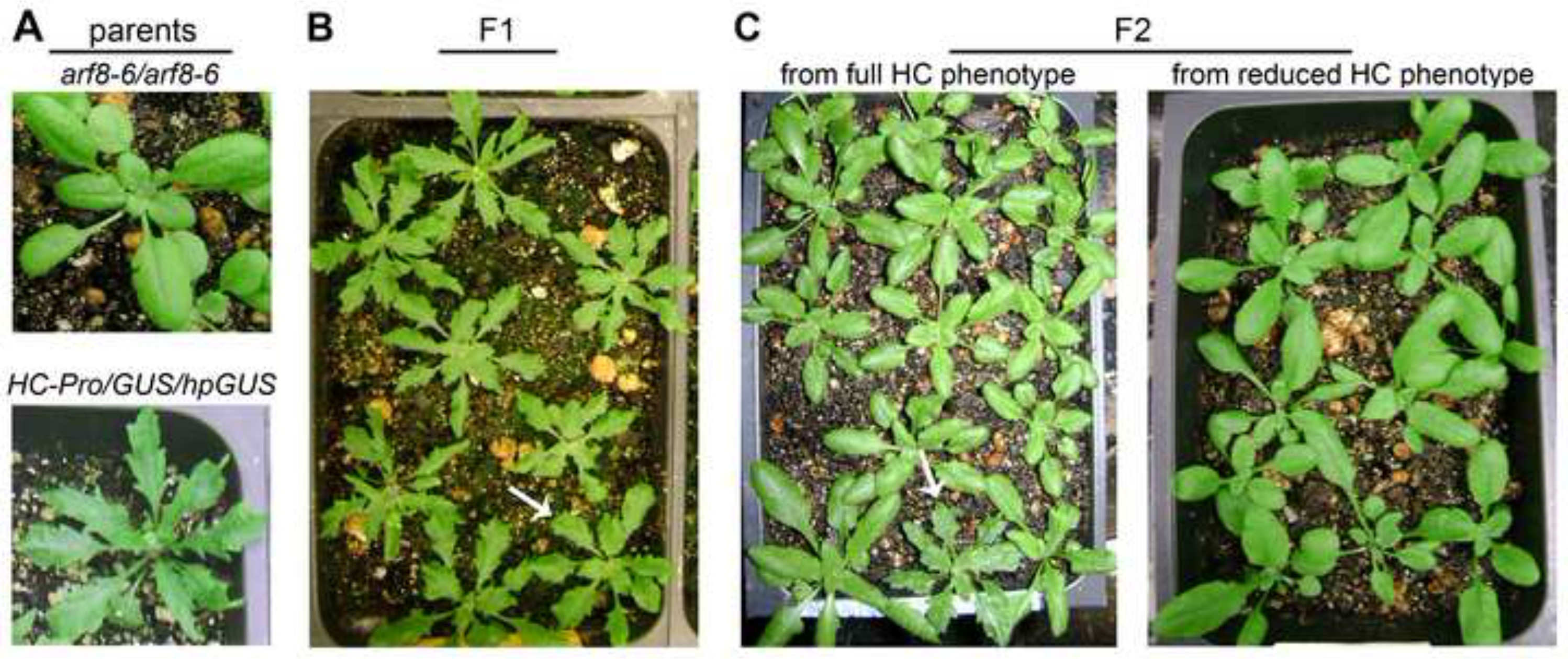
The P1/HC-Pro phenotype is largely retained in the F1 but lost in the F2 generation of a cross with the *arf8-*6 mutant. Representative parental plants plus F1 and F2 progeny of the cross between *arf8-6* and P1/HC-Pro lines are shown. **(A)** The *arf8-6/arf8-6* parental line has a wild-type phenotype, whereas the *P1/HC-Pro*/GUS/hpGUS parent exhibits the characteristic phenotype of our *P1/HC-Pro* lines. **(B)** P1/HC-Pro phenotypic F1 progeny of the cross, showing that most have the full P1/HC-Pro phenotype and that only a minority have a reduced phenotype (arrow). **(C)** Full and reduced P1/HC-Pro phenotype F1 plants from which RNA was isolated were allowed to self. Seeds from each of the F1 phenotypic groups were pooled separately before planting. No P1/HC-Pro phenotypic F2 progeny carrying *arf8-6* were obtained from the reduced phenotype F1 plants (right pot), while a minority of the F2 progeny from the full phenotype F1 plants exhibited the P1/HC-Pro phenotype (left pot, arrow).

### The P1/HC-Pro phenotype is largely lost in the F2 generation of the *arf8-6* cross, but the loss is independent of whether the *arf8-6* mutation is hemizygous or homozygous

Because an effect of the *arf8-6* mutation on the phenotype of our *P1/HC-Pro* line might be more evident when plants are homozygous rather than hemizygous for the mutation, we examined the phenotypes of the F2 progeny of the cross (after self-pollination of the F1 plants). Progeny of the slightly reduced P1/HC-Pro phenotypic F1 plants had completely lost the phenotype in the F2 generation (Fig 1, F2) whether plants were hemizygous or homozygous for *arf8-6*. P1/HC-Pro phenotypic progeny obtained from the slightly reduced P1/HC-Pro phenotype F1 plants all lacked the *arf8-6* T-DNA insertion. Thus, although complete loss of P1/HC-Pro phenotype occurred in this population of the F2 progeny of the *arf8-6* cross, the loss did not require that *arf8-6* be homozygous. The P1/HC-Pro phenotype was also lost in the F2 progeny of the fully P1/HC-Pro phenotypic F1 plants, but was not total in this case (Fig 1, F2). Only about 25% (67/271) of this F2 population had a P1/HC-Pro phenotype, and most (50/67) had a reduced phenotype. All of these reduced P1/HC-Pro phenotype plants had *arf8-6*, but only 11 out of the 50 were homozygous for the mutation. Thus, progressive loss of P1/HC-Pro phenotype occurred in successive generations of the *arf8-6* cross, but was independent of whether the *arf8-6* mutation was hemizygous or homozygous. Of the 17 fully P1/HC-Pro phenotypic plants obtained in this F2 population, 10 had the *arf8-6* mutation. Two of the 10 were homozygous for *arf8-6*, showing that the full P1/HC-Pro phenotype can occur even in the complete absence of *ARF8* expression.

Altogether, our results suggest that some mechanism other than reduced *ARF8* expression is responsible for alleviation and loss of P1/HC-Pro phenotype in plants carrying the *arf8-6* mutation. Because the *arf8-6* mutation is a SALK line T-DNA insertion mutation, one likely candidate is transcriptional silencing induced by the T-DNA. The T-DNAs used to generate SALK lines and several other T-DNA insertion lines carry an extraneous CaMV 35S promoter that is known to transcriptionally silence 35S promoter-driven transgenes elsewhere in the plant genome [15].

Such unexpected silencing effects can generate misleading results, especially in studies involving viral suppressors of silencing because such suppressors are not effective against transcriptional silencing. The *P1/HC-Pro* transgene in our lines and the one in the work of Jay et al. (2011) are both expressed from the 35S promoter. Therefore, one likely possibility is that transcriptional silencing is the mechanism responsible for alleviation and loss of P1/HC-Pro phenotype in plants also carrying the *arf8-6* mutation.

### Indications of 35S promoter-induced transcriptional silencing are evident in the reduced P1/HC-Pro phenotype F1 progeny of the *arf8-6* cross

To see whether transcriptional silencing of 35S promoter-driven transgenes was occurring in our *arf8-6* mutant plants, we performed RNA gel blot analysis of high and low molecular weight RNA isolated from F1 and F2 progeny of the *arf8-6* cross. For the purpose of the molecular analysis, the *P1/HC-Pro* line we used for the cross contained two additional transgenes that provide an assay for silencing suppression [14]. These additional loci are a sense *uidA* transgene (*GUS*) encoding β-glucuronidase and a hairpin transgene (*hpGUS*) that post-transcriptionally silences the *GUS* locus [16]. Expression of each of the three transgenes in this *P1/HC-Pro* line is under the control of the Cauliflower Mosaic Virus (CaMV) 35S-promoter. The *hpGUS* and *GUS* transgenes function not only as reporters for suppression of post-transcriptional silencing by P1/HC-Pro, but also as additional indicators of transcriptional silencing because each is expressed from a 35S promoter. On their own, however, these two transgenes do not induce transcriptional silencing [14, 16].

For the RNA gel blot analysis, we examined RNA from plants that had *P1/HC-Pro*, the *arf8-6* mutation, as well as both the *hpGUS* and *GUS* transgenes. In the F1 progeny of the cross, 50 plants were heterozygous for the *arf8-6* mutation and contained all three of the other transgenes. Of these, 38 plants retained the full P1/HC-Pro phenotype, while 12 showed a slight reduction in the phenotype. RNA gel blot analysis shows that fully P1/HC-Pro phenotypic *arf8-6* plants have the same levels as the *P1/HC-Pro* parent for all high and low molecular weight RNA species examined (Fig 2, lanes 1-6): *P1/HC-Pro* is expressed at about the same level and suppresses post-transcriptional silencing equally well, as indicated by the highly increased accumulation of *GUS* and *hpGUS* mRNA compared to the non-*P1/HC-Pro* controls (Fig 2, compare lanes 1-6 with lanes 15-19). The *GUS* locus gives rise to a transcript that is slightly longer than the full-length *hpGUS* transcript; however, a prominent high molecular weight *hpGUS* RNA species that also accumulates corresponds to the loop of the hairpin [14]. With respect to low molecular weight RNA species, the fully P1/HC-Pro phenotypic *arf8-6* plants and the *P1/HC-Pro* parent show the same increased accumulation of miRNAs and of primary siRNAs, which are the siRNAs from the stem of the *hpGUS* transcript (Fig 2, compare lanes 1-6 with lanes 15-19).

**Fig 2.**
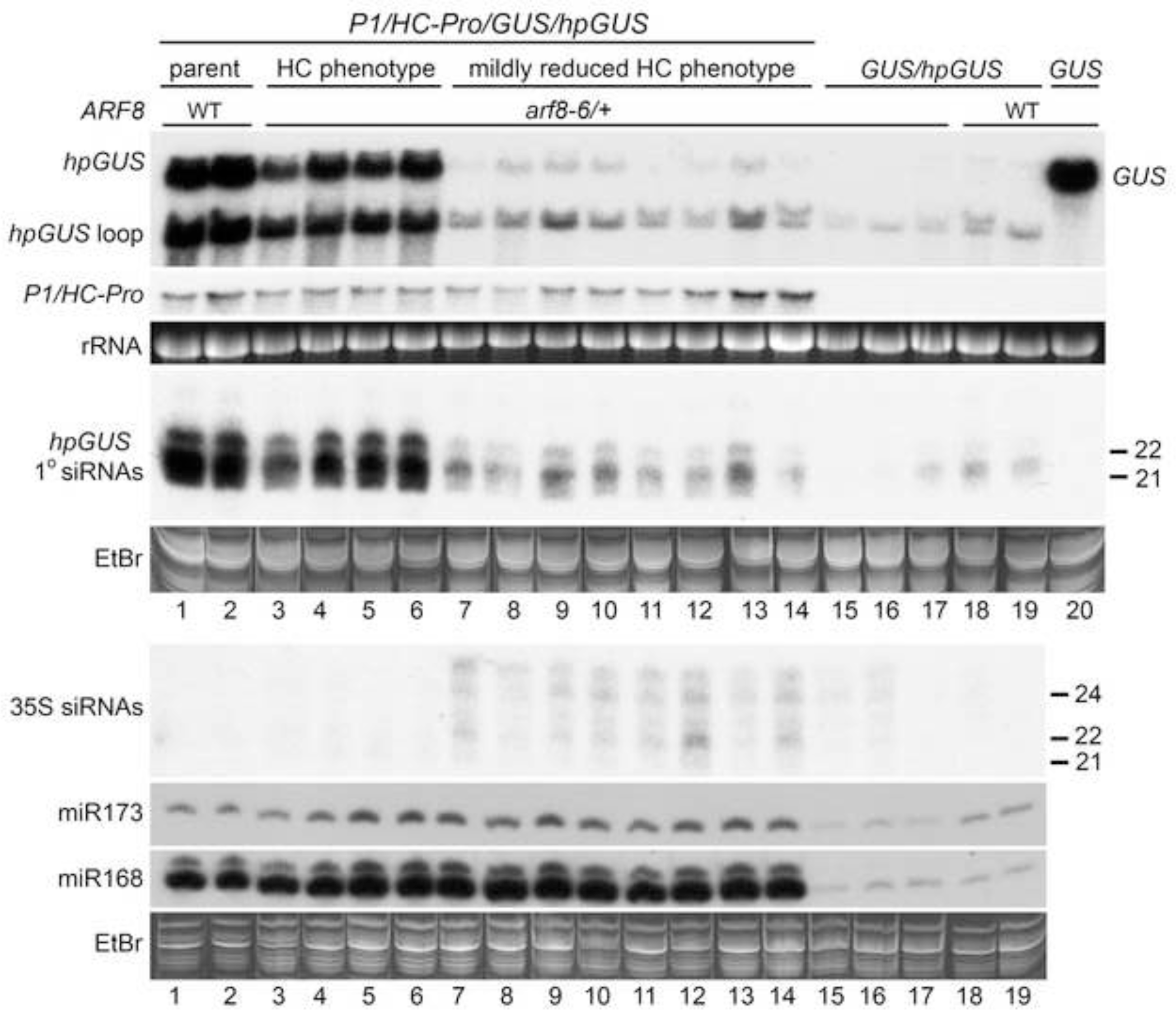
Transcriptional silencing is evident in F1 progeny of the *arf8-6* cross that have a reduced P1/HC-Pro phenotype. Accumulation of high and low molecular weight RNA from parental and F1 progeny plants having the indicated genotypes and phenotypes was determined using RNA gel blot analysis with RNA probes. RNA from the *GUS* line was included as a control to show the position of the *GUS* transcript. Grouped lanes are all from the same gel. The high molecular weight blot was hybridized with a probe for *GUS* and *hpGUS*, then stripped and probed for *P1/HC-Pro*. Two separate low molecular weight (LMW) RNA gels were run. One LMW blot was probed for *hpGUS* primary siRNAs, while the other was probed for 35S siRNAs, then successively stripped and probed for the indicated miRNAs. Ethidium bromide (EtBr) stained rRNA and the major RNA species in LMW RNA are shown as loading controls.

Although the difference in phenotype between F1 progeny having the full P1/HC-Pro phenotype and those having a slightly reduced phenotype is quite subtle, the difference in the RNA data of these two groups is very striking. The RNA data for the slightly reduced phenotype plants present a very complicated picture because transcriptional silencing and P1/HC-Pro suppression of post-transcriptional silencing both appear to be occurring to some degree. 35S promoter siRNAs are associated with transcriptional silencing induced by T-DNA insertion mutants [17], providing a good indicator for 35S promoter homology-dependent transcriptional silencing. 35S siRNAs were not evident in any of the fully P1/HC-Pro phenotypic *arf8-6* F1 plants (Fig 2, lanes 3-6) or in plants that lacked the *arf8-6* mutation (Fig 2, lanes 1-2 and 18-19). In contrast, we detected 35S promoter siRNAs in RNA from all of the slightly reduced P1/HC-Pro phenotype *arf8-6* F1 plants assayed (Fig 2, lanes 7-14), indicating that transcriptional silencing had begun in this population. Consistent with the very subtle reduction in phenotype, however, accumulation of P1/HC-Pro mRNA appears unaffected (Fig 2, compare lanes 7-14 with lanes 1-6). In contrast, accumulation of *GUS* and *hpGUS* mRNA as well as primary siRNAs from *hpGUS* is much lower than in the fully P1/HC-Pro phenotypic plants (Fig 2, compare lanes 7-14 to lanes 3-6), although still higher in some cases than in plants lacking *P1/HC-Pro* (Fig 2 compare lanes 7-14 to 15-19). Thus, despite the onset of transcriptional silencing, P1/HC-Pro suppression of post-transcriptional silencing appears to be occurring in at least some of the mildly reduced phenotype F1 progeny. P1/HC-Pro activity in the mildly reduced phenotype F1 progeny is more clearly demonstrated by an increase in miRNA accumulation comparable to that in fully P1/HC-Pro phenotypic plants (Fig 2 compare lanes 7-14 to 1-6 and 15-19).

Thus, the *P1/HC-Pro/arf8-6* F1 plants are a mixed population. Most are completely normal for transgene expression and P1/HC-Pro activity, but some have started to transcriptionally silence 35S promoter-driven transgenes. It is interesting to note that 35S promoter-induced transcriptional silencing appears to have started even in *arf8-6* progeny that do not have the *P1/HC-Pro* transgene: 35S promoter siRNAs are very faintly detectable in RNA from two of the three *arf8-6/GUS/hpGUS* plants we analyzed (Fig 2, compare lanes 15-16 with lanes 18-19).

### 35S promoter-induced transcriptional silencing is widespread and well developed in the F2 progeny of the *arf8-6* cross

RNA gel blot analysis of the F2 progeny of the *arf8-6* cross confirms that transcriptional silencing of 35S promoter-driven transgenes is occurring, even in the absence of the *P1/HC-Pro* transgene. All three of the *arf8-6^-/-^/GUS/hpGUS* plants analyzed are transcriptionally silenced for the *GUS* and *hpGUS* transgenes. These plants show high levels of 35S siRNAs and have no detectable *GUS* and *hpGUS* mRNA or primary siRNAs from *hpGUS*, in contrast to *GUS/hpGUS* controls lacking *arf8-6* (Fig 3, compare lanes 15-17 with lanes 18-19). Similarly, *P1/HC-Pro/arf8-6^-/-^/GUS/hpGUS* F2 plants that have completely lost the P1/HC-Pro phenotype are clearly transcriptionally silenced for *P1/HC-Pro* as well as for the *GUS* and *hpGUS* transgenes. These plants have easily detectable levels of 35S siRNAs and accumulate little or no P1/HC-Pro mRNA and no *GUS* and *hpGUS* mRNA or primary siRNAs from *hpGUS*, compared to *P1/HC-Pro* controls lacking the *arf8-6* mutation (Fig 3, compare lanes 7-14 with lanes 1-2). Surprisingly, however, miR168 accumulation is slightly elevated compared to *non-P1/HC-Pro* controls (Fig 3, compare lanes 7-14 with lanes 15-19).

**Fig 3.**
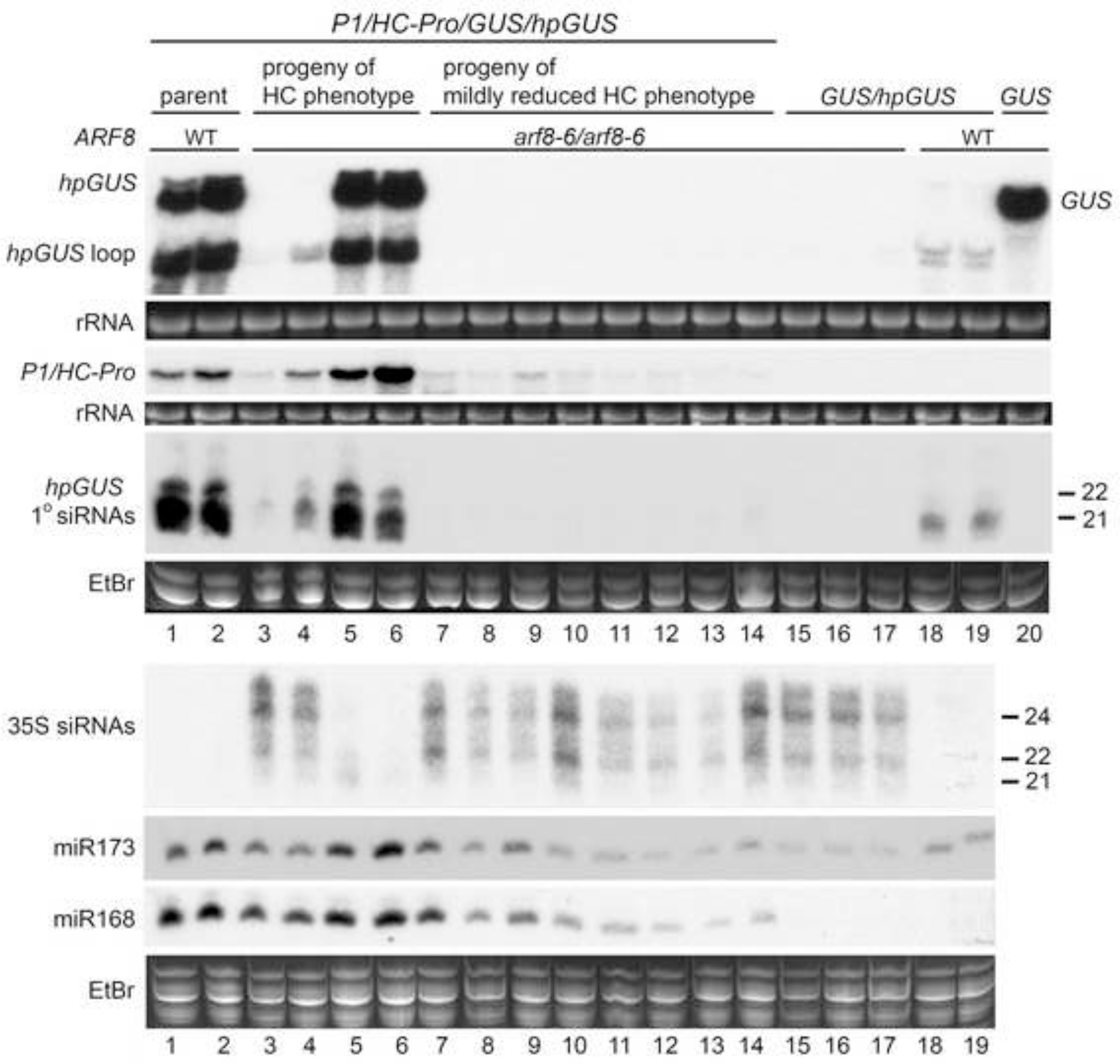
Transcriptional silencing is widespread in the F2 progeny of the *arf8-6* cross. Accumulation of high and low molecular weight RNA from the original parental lines and from F2 progeny having the indicated genotypes and F1 parental phenotypes was determined using RNA gel blot analysis with RNA probes. RNA from the *GUS* line was included as a control to show the position of the *GUS* transcript. Grouped lanes are all from the same gel. Two high molecular weight (HMW) RNA gels were run. One HMW blot was hybridized with a probe for *GUS* and *hpGUS*, while the other was probed for *P1/HC-Pro*. Two low molecular weight (LMW) RNA gels were run. One LMW blot was probed for *hpGUS* primary siRNAs, while the other was probed for 35S siRNAs, then successively stripped and probed for the indicated miRNAs. Ethidium bromide (EtBr) stained rRNA and the major RNA species in LMW RNA are shown as loading controls.

As discussed above, some *P1/HC-Pro/arf8-6^-/-^F2* plants had a P1/HC-Pro phenotype (Fig 1, F2). Most of these had a reduced phenotype, but a few had the full P1/HC-Pro phenotype. RNA gel blot analysis shows that 35S promoter-induced transcriptional silencing had started in the reduced phenotype plants, as evidenced by the presence of 35S siRNAs as well as reduced levels of *P1/HC-Pro, GUS* and *hpGUS* mRNA and *hpGUS* primary siRNAs (Fig 3,compare lanes 3-4 with lanes 1-2). In contrast, 35S promoter-induced transcriptional silencing does not appear to have started in the few *P1/HC-Pro/arf8-6^-/-^F2* plants that still have the full P1/HC-Pro phenotype. These plants look identical to the *P1/HC-Pro* parent for all high and low molecular weight RNA species examined (Fig 3, compare lanes 5-6 with lanes 1-2). Thus, 35S promoter-induced transcriptional silencing appears well established in the vast majority of F2 progeny of the *arf8-6* cross, supporting our hypothesis that transcriptional silencing is the mechanism responsible for loss of P1/HC-Pro phenotype in *P1/HC-Pro/arf8-6* plants. The few *P1/HC-Pro/arf8-6* plants that retain some degree of P1/HC-Pro phenotype are ones in which transcriptional silencing has not yet been completely established.

### The *arf8-8* loss of function point mutation does not alleviate P1/HC-Pro-mediated developmental defects

To test further whether loss of *ARF8* function can alleviate the P1/HC-Pro phenotype, we used the *arf8-8* mutant line, which has a point mutation at the junction of the 5^th^ intron and 6^th^ exon of *ARF8* [18]. The *arf8-8* mutation causes vegetative and flower growth phenotypes indistinguishable from those caused by the loss-of-function T-DNA insertion mutation *arf8-3*, both as a single mutant and in combination with the *ARF6* loss-of-function mutation, *arf6-2* (Fig 4). Because the *ARF8* and *ARF6* genes act partially redundantly [19], the loss of function of either gene alone has only a subtle effect, whereas the double mutant is profoundly affected (Fig 4 and [19]). Thus, *arf8-8* appears phenotypically as a null allele. We crossed our *P1/HC-Pro/GUS/hpGUS* line to the *arf8-8* mutant line and examined the phenotypes of the F1 progeny. There were two classes of phenotype: P1/HC-Pro and wild type. All of the F1 progeny we obtained that had *P1/HC-Pro* in the heterozygous *arf8-8* background retained the distinctive P1/HC-Pro phenotype (Fig 5, F1). None of the non-phenotypic plants had the *P1/HC-Pro* transgene, and none of the *P1/HC-Pro* heterozygous *arf8-8* plants showed any reduction in P1/HC-Pro phenotype. The P1/HC-Pro phenotype was also retained in the F2 generation of the cross, whether plants were homozygous or heterozygous for the *arf8-8* mutation (Fig 5, F2).

**Fig 4.**
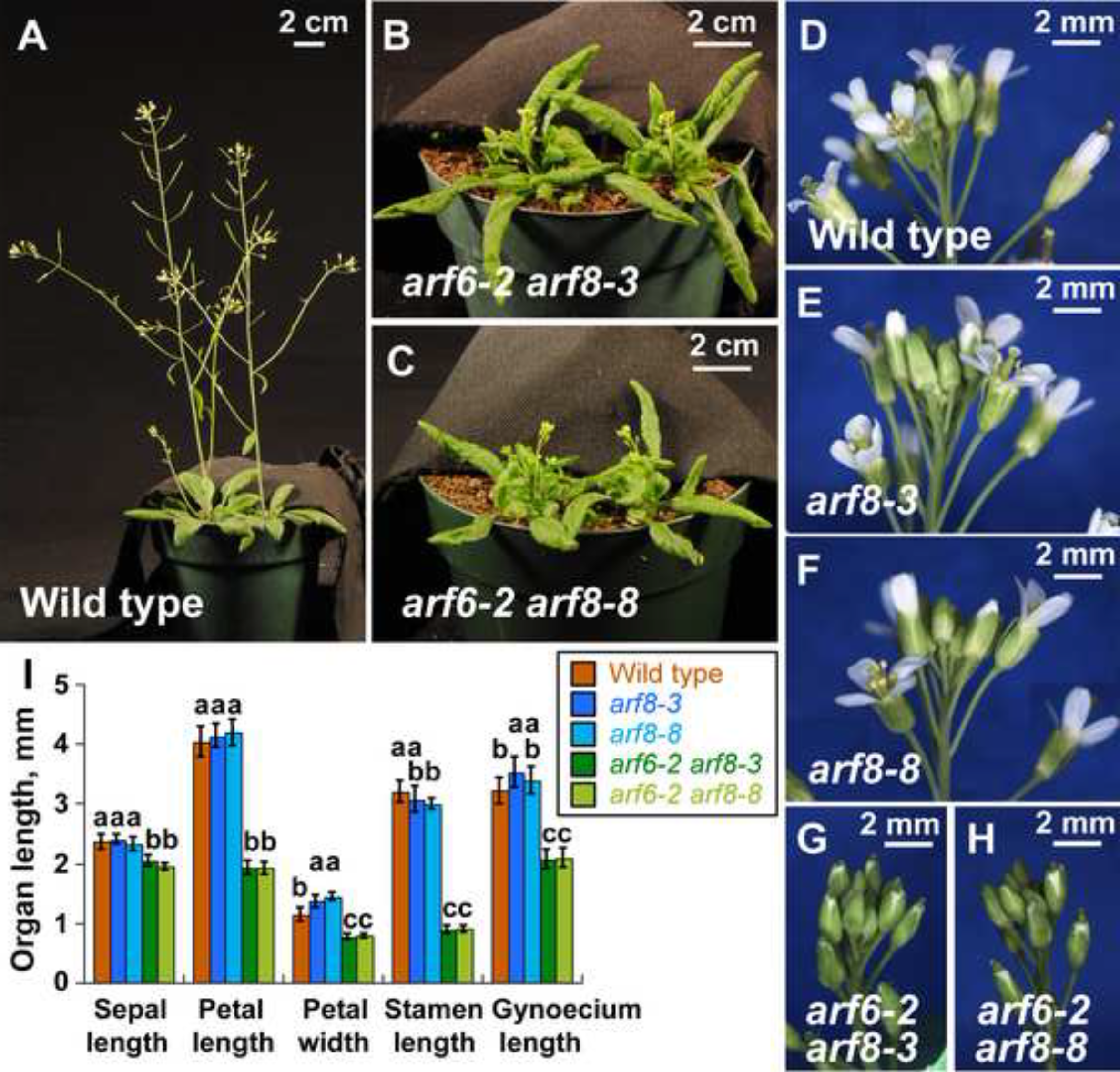
Comparison of *arf8-3* and *arf8-8* developmental phenotypes. **(A-C)** 5-week-old wild-type, *arf6-2 arf8-3*, and *arf6-2 arf8-8* plants. **(D-H)** Inflorescence apices of 5-week-old plants of indicated genotypes. Panels D, E, and F were each assembled from multiple overlapping photographs**. (I)** Measurements of floral organ lengths and petal widths of mature flowers of each genotype. Data are means α sd (n=13 to 18 flowers). In I, letters above each bar indicate values that were not statistically distinguishable by ANOVA at P<0.05 based on Tukey's HSD statistic, calculated within each measurement class. As ARF6 and ARF8 act redundantly, the *arf6-2* background exposes the full phenotypic effects of the *arf8* mutations.

**Fig 5.**
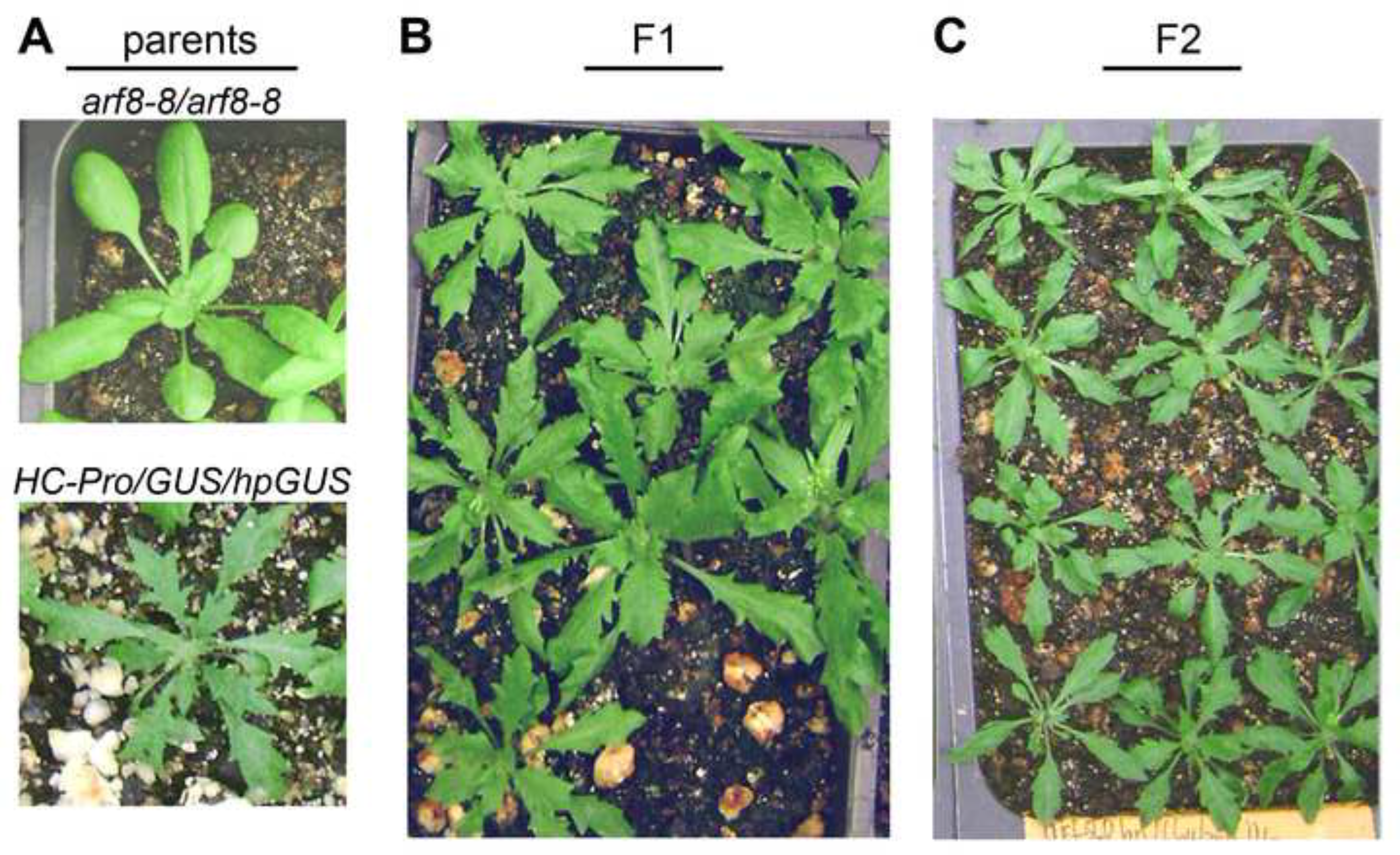
The *arf8-8* mutation has no effect on the P1/HC-Pro phenotype in either the F1 or F2 generation of a cross to our *P1/HC-Pro* line. Representative parental plants plus F1 and F2 progeny of the cross between *arf8-8* and *P1/HC-Pro* lines are shown. **(A)** The *arf8-8/arf8-8* parental line has a wild-type phenotype, whereas the *P1/HC-Pro/GUS/hpGUS* parent exhibits the characteristic phenotype of our *P1/HC-Pro* lines. **(B)** P1/HC-Pro phenotypic F1 progeny of the cross, showing that all have the full P1/HC-Pro phenotype. **(C)** P1/HC-Pro phenotypic F1 plants from which RNA was isolated were allowed to self, and seeds were pooled before planting. All *P1/HC-Pro* F2 progeny carrying arf8-8 exhibited the full P1/HC-Pro phenotype.

### The *arf8-8* loss of function point mutation does not affect P1/HC-Pro function

Because P1/HC-Pro phenotype was unaffected by the *arf8-8* mutation (Fig 5), we expected that molecular measures of P1/HC-Pro function would also be unaffected. RNA gel blot analysis of high and low molecular weight RNA isolated from F1 and F2 progeny of the *arf8-8* cross support that expectation. *P1/HC-Pro* plants hemizygous (Fig 6) or homozygous (Fig 7) for *arf8-8* look nearly identical to the *P1/HC-Pro* parent for all high and low molecular weight RNA species examined (Fig 6 compare lanes 3-7 with lanes 1-2; Fig 7 compare lanes 3-6 with lanes 1-2). P1/HC-Pro suppresses the *hpGUS* transgene-induced silencing in these plants, as indicated by the highly increased accumulation of *GUS* and hpGUS mRNA, and mediates increased accumulation of miRNAs and hpGUS primary siRNAs compared to the *non-P1/HC-Pro* controls (Fig 6, compare lanes 3-7 and 1-2 with lanes 8-14; Fig 7 compare lanes 3-6 and 1-2 with lanes 7-12). Thus, reduced levels of functional *ARF8* in the absence of 35S promoter-induced transcriptional silencing affect neither P1/HC-Pro phenotype nor molecular measures of P1/HC-Pro function.

**Fig 6.**
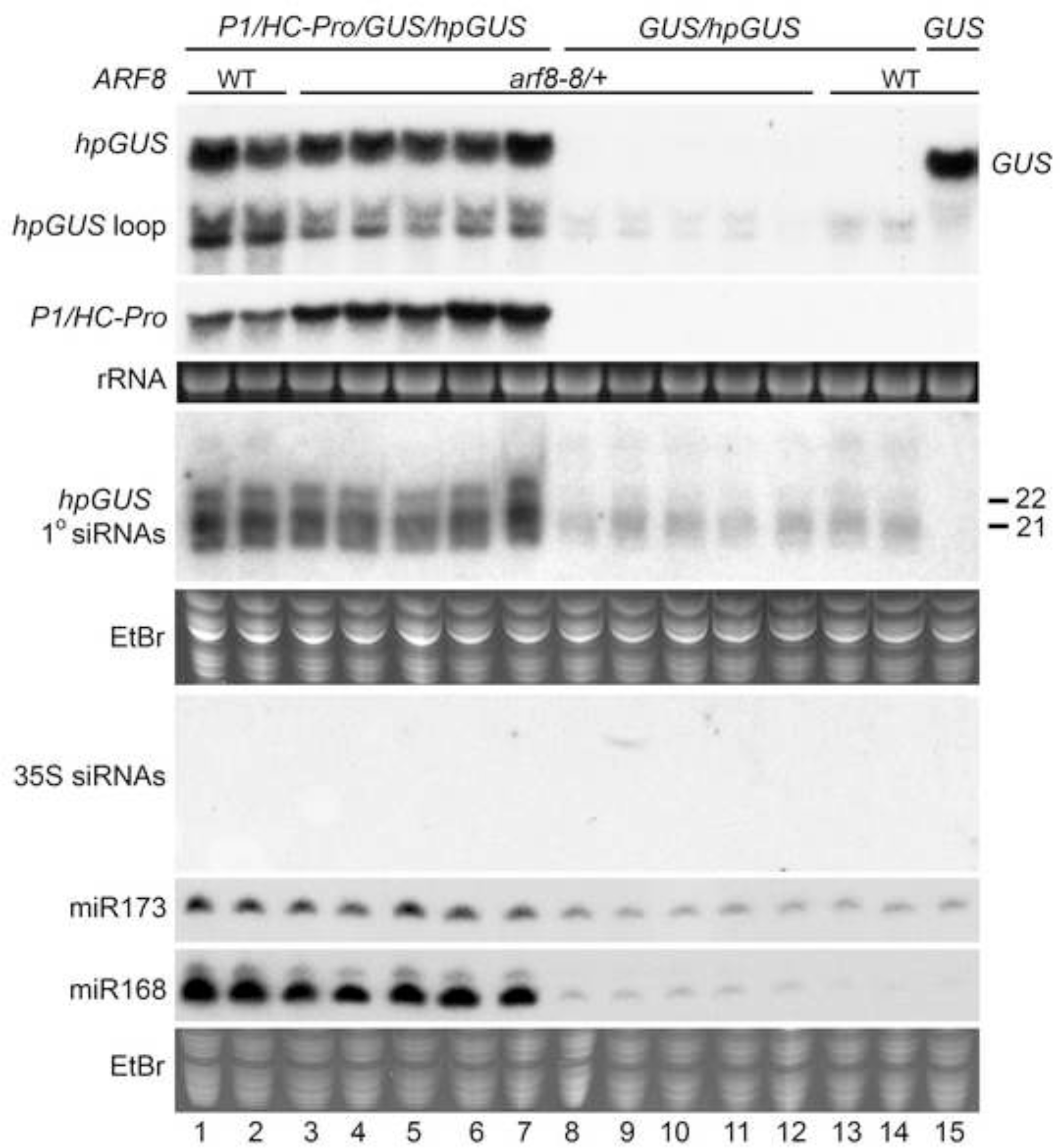
The *arf8-8* mutation has no effect on molecular measures of P1/HC-Pro function in the F1 generation of a cross to our *P1/HC-Pro* line. Accumulation of high and low molecular weight RNA from parental and F1 progeny plants having the indicated genotypes was determined using RNA gel blot analysis with RNA probes. RNA from the *GUS* line was included as a control to show the position of the *GUS* transcript. Grouped lanes are all from the same gel. The high molecular weight (HMW) blot was hybridized with a probe for *GUS* and *hpGUS*, then stripped and probed for *P1/HC-Pro*. Two separate low molecular weight (LMW) RNA gels were run. One LMW blot was probed for *hpGUS* primary siRNAs, while the other was probed for 35S siRNAs, then successively stripped and probed for the indicated miRNAs. Ethidium bromide (EtBr) stained rRNA and the major RNA species in LMW RNA are shown as loading controls.

**Fig 7.**
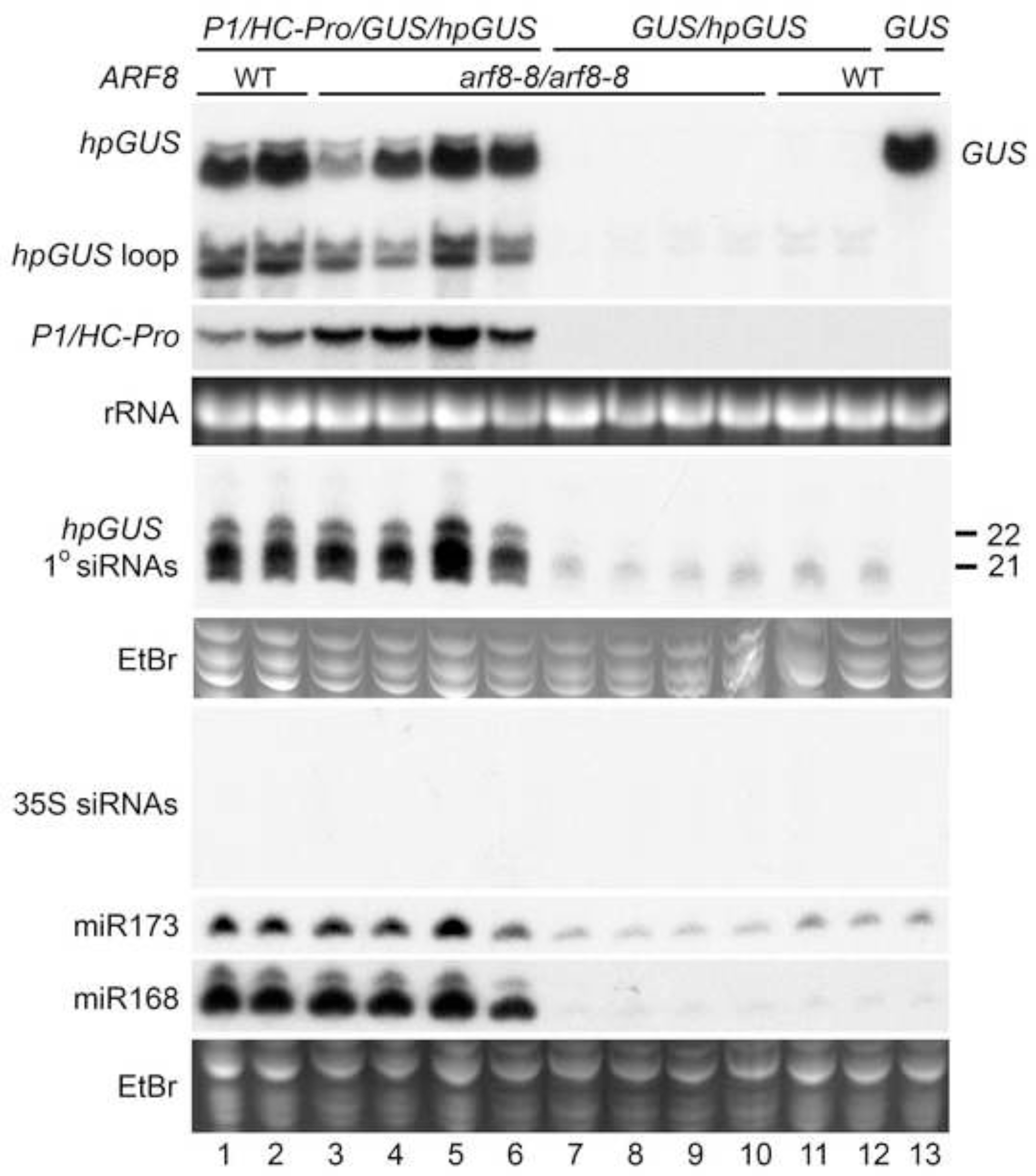
The arf8-8 mutation has no effect on molecular measures of P1/HC-Pro function in the F2 generation of a cross to our *P1/HC-Pro* line. Accumulation of high and low molecular weight RNA from parental and F2 progeny plants having the indicated genotypes was determined using RNA gel blot analysis with RNA probes. RNA from the *GUS* line was included as a control to show the position of the *GUS* transcript. Grouped lanes are all from the same gel. Two high molecular weight (HMW) RNA gels were run. One HMW blot was hybridized with a probe for *GUS* and *hpGUS*, while the other was probed for *P1/HC-Pro*. Two low molecular weight (LMW) RNA gels were run. One LMW blot was probed for *hpGUS* primary siRNAs, while the other was probed for 35S siRNAs, then successively stripped and probed for the indicated miRNAs. Ethidium bromide (EtBr) stained rRNA and the major RNA species in LMW RNA are shown as loading controls.

## Discussion

Our results clearly show that *ARF8* is not the key factor responsible for developmental defects in *Arabidopsis* expressing a TuMV *P1/HC-Pro* transgene. Moderate upregulation of *ARF8* was originally reported to underlie developmental defects in *Arabidopsis* induced by P1/HC-Pro and two other viral suppressors of silencing [11]. The identification of *ARF8* as a candidate for this role came from determining what miRNA targets are upregulated in common in *Arabidopsis* expressing any one of the viral suppressors, P1/HC-Pro, P19, and P15, as well as in *Arabidopsis hen1-1* and *dcl1-9* mutants. The upregulation in all cases was about 1.5-to 2-fold suggesting that loss of just one copy of *ARF8* would greatly reduce or eliminate the developmental defects caused by the viral suppressors. Alleviation of the developmental defects by one copy of the *arf8-6* SALK insertion mutation was then taken as evidence that misregulation of *ARF8* was the key factor underlying viral suppressor-mediated developmental anomalies.

We were led to question the conclusions of the Voinnet group [11] for three reasons. The first was our prior work, which showed no correlation between P1/HC-Pro phenotype and *ARF8* expression [9]. The second reason was a paradoxical result presented in the gel blot analysis of RNA isolated from *P1/HC-Pro* plants that carried the *arf8-6* mutation and a hairpin transgene that targets the gene encoding chalcone synthase (CHS) [[11]: Fig 5E]. The paradoxical result was that the gel blot analysis showed a total absence of *CHS* siRNAs in those plants, although that same panel of the figure presented evidence that P1/HC-Pro was suppressing the hairpin transgene-induced post-transcriptional silencing. It has been well established, however, that primary siRNAs, such as those that would derive from the stem of a hairpin transgene, are not eliminated in P1/HC-Pro suppression of silencing [13, 14, 20, 21]. Therefore, *CHS* siRNAs should have been present if P1/HC-Pro suppression of silencing was truly occurring, and their absence strongly suggested that some confounding factor had affected the Jay et al. P1/HC-Pro studies [11]. Lastly, the P1/HC-Pro phenotypes differ in several aspects from those caused by loss of miR167 regulation of *ARF6* or *ARF8* [12].

Our work clearly shows that all the effects of the *arf8-6* T-DNA insertion mutation on P1/HC-Pro phenotype and on molecular indicators of P1/HC-Pro function are due to 35S promoter homology-dependent transcriptional silencing induced by the *arf8-6* T-DNA insertion. In addition, the *arf8-8* loss of function point mutation has no effect on P1/HC-Pro phenotypes or on molecular measures of P1/HC-Pro function, further arguing against *ARF8* as the key factor responsible for developmental defects caused by *P1/HC-Pro* expression. Thus, the exact mechanism responsible for P1/HC-Pro-mediated developmental defects remains an open question. Interference with miRNA-controlled developmental pathways is a very attractive hypothesis, but ectopic overexpression of *DCL1* has been shown to separate P1/HC-Pro phenotype from effects on general miRNA biogenesis and function [9]. Therefore, one or a few key miRNA-controlled factors might, in fact, underlie the developmental defects caused by P1/HC-Pro; however, our work shows that *ARF8* is not one of these key factors.

## Materials and Methods

### Arabidopsis lines

The mutant and transgenic lines used are all in the Columbia (Col-0) ecotype and have been described in previous studies. The *P1/HC-Pro/GUS/hpGUS* line is hemizygous for the *P1/HC-Pro* transgene and was previously published as *P1/HC-Pro/6b4/306* [14]. The *arf8-6* line is the University of Wisconsin T-DNA line WiscDsLox324F09 [22]. The *arf8-8* line has an ethyl methane sulfonate (EMS)-generated G to A mutation in a splice acceptor site in *ARF8* [18].

### Genotyping

Hemizygous and homozygous *arf8-8* plants were genotyped using the PCR primers and EcoNI digestion of the PCR product as described previously [18]. For hemizygous and homozygous *arf8-6* genotyping, the T-DNA primer LB (5′-AACGTCCGCAATGTGTTATTAAGTTGTC-3′) was used with primers ARF8-6LP (5′-CGAGGAAAGGTGAAACCTAC-3′) and ARF8-6RP(5′-AGCTGTCAACATCTGGATTGG-3′). Primers ARF8-6LP and ARF8-6RP amplify a 1014 bp product from the wild-type locus, and primers ARF8-6RP and LB amplify a 500 bp product from the *arf8-6* locus. Genotyping for the *hpGUS* (= 306) and *GUS* (= 6b4) transgenes was performed as described previously [14]. *P1/HC-Pro* primers HC-F (GTGCCCAGAAGTTCAAGAGC) and HC-R (GTCAACGACTATGCCACTCCAACC) were used to confirm the presence of the *P1/HC-Pro* transgene in combination with phenotypic selection. P1/HC-Pro phenotype was evaluated based on the degree of dwarfing and leaf serration compared to wild-type *Arabidopsis*.

### RNA isolation and gel-blot analysis

High and low molecular weight total RNA was isolated from aerial tissues of individual flowering plants and gel blot analysis performed as described previously [9, 14] [α-^32^P]UTP-labeled antisense RNA probes to detect *GUS, hpGus*, and P1/HCPro mRNAs or the sense RNA probes to detect *hpGUS* primary siRNAs and 35S-promoter siRNAs were prepared using an Ambion MAXIscript *in vitro* transcription kit (Ambion, http://www.ambion.com) as described previously [14, 17]. The probes were hybridized to mRNA blots at 68°C in Ambion ULTRAhyb buffer, or to siRNA blots at 42°C in Ambion ULTRAhyb-oligo buffer. Antisense oligonucleotide probes for detection of miRNAs were prepared by end-labeling with ^32^P using T4 polynucleotide kinase (New England Biolabs) and hybridized to miRNA blots at 42°C in Ambion ULTRAhyb-oligo buffer.

### *arf* mutant phenotypic analyses

Flower organs were dissected from mature open flowers of wild-type and *arf8* single mutants (2-3 flowers per apex), or from arrested flower buds at the equivalent position on the inflorescence of *arf6 arf8* double mutants (5-7 flowers per apex). Organs were placed flat on agar plates, and measured using a camera lucida attached to a dissecting microscope [18]. Mean lengths or widths measured from at least two sepals, petals, and long stamens for each flower were used in the summary graph in Figure 4.

